# Dual role of the OMM E3 Ub ligase MARCH5 in *de novo* peroxisome biogenesis and mitochondrial quality control through direct regulation of Pex26

**DOI:** 10.64898/2026.05.06.723346

**Authors:** Debduti Bhattacharjee, Claudia Christiane Bippes, Guiling Zhao, Liron Boyman, Mehari M. Weldemariam, Maureen A. Kane, Albert Neutzner, Mariusz Karbowski

## Abstract

Recent evidence indicates that mitochondria, through the activity of the E3 Ub ligase MARCH5, are critical for *de novo* peroxisome biogenesis. Here we report that peroxisome biogenesis factor Pex26 is a MARCH5 client protein. In peroxisome-containing cells, MARCH5 interacts with Pex26 and facilitates the transfer of newly synthesized Pex26 from the OMM to peroxisomes. MARCH5 also controls peroxisomal delivery of other candidate peroxins in peroxisome-containing cells. On the other hand, in peroxisome-deficient cells, the turnover rate of Pex26 is dramatically increased, and MARCH5 targets this protein for p97-dependent proteasomal degradation. Both activities are mediated by MARCH5-dependent Pex26 ubiquitination. Knockout of Pex26 induces the accumulation of cells containing Tom20-positive, Catalase-deficient pre-peroxisomes. Further supporting the critical role of MARCH5 in peroxisome biogenesis, these structures are absent in Pex26/MARCH5 double knockout cells. The data support the model, where in peroxisome-containing cells, MARCH5 acts as a peroxisome biogenesis factor, while with defective peroxisome biogenesis, as in Zellweger syndrome cells, it protects mitochondria from potentially toxic accumulation of peroxins on the OMM.

## Introduction

The interplay between mitochondria and peroxisomes is critical for lipid metabolism, regulation of reactive oxygen species (ROS), and ATP generation. Peroxisomes form by the growth and division of pre-existing organelles but can also emerge *de novo.* In *de novo* biogenesis pathway peroxisomes are thought to arise from the ER^1^. However, extending or disputing the ER-centric view of *de novo* peroxisome biogenesis that implicated ER as a sole platform for *de novo* peroxisome biogenesis, recent works show that in this pathway, a subset of pre-peroxisomes develops from the mitochondria, and some peroxisomal biogenesis factors, including Pex3 and Pex14, could have functions in this process^2–4^. Furthermore, many proteins originally discovered to be mitochondrial, including dynamin-related protein 1 (Drp1)^5^, mitochondrial fission factor (Mff)^5^, Fis1^5^ and GTPase Mfn2^6^, E3 ubiquitin (Ub) ligase MARCH5^3,4,7,8^, deubiquitinase Usp30^9–11^, and mitophagy factor BNIP3L/NIX^12^, localize to and control peroxisomes, while others like innate immunity factor MAVS^13^ use mitochondria and peroxisomes as signaling platforms. Some luminal peroxisomal proteins are likely to have bacterial/mitochondrial origin^14^. The presence of shared proteins suggests an evolutionary link where proteins initially targeted to mitochondria were later recruited to peroxisomes^14^. The possibility that mitochondria are a critical platform in peroxisome biogenesis is supported by recent reports ^2–4^.

Grounded on the dominant view that ER is the major platform in *de novo* peroxisome biogenesis^15–19^ it has been proposed that mitochondrial localization of peroxins in peroxisome-deficiency models represent mistargeting of peroxisomal proteins occurring in the absence of peroxisomes. However, another interpretation is that mitochondria play a role in peroxisomal biogenesis ^2–4^. The report showing that protein phosphatase type 2C (Ptc5) transfers through the mitochondria *en route* to its destination in peroxisomes^20^, could serve as proof of principle data, supporting this underestimated possibility. Mitochondrial steps in *de novo* peroxisome biogenesis require the activity of OMM integral E3 Ub ligase MARCH5 ^3,4^.

MARCH5 has many mitochondrial functions, including regulation of mitochondrial fusion and fission^7,21,22^, removal of exogenous proteins from the OMM^23^, control of mitochondrial protein import fidelity^24^, and regulation of mitochondrial steps in apoptosis^21,25,26^. MARCH5 also localizes to peroxisomes^3,4,8^, and controls biogenesis of these organelles, via a currently unknown mechanism. Like MARCH5, other primarily mitochondrial components of the Ub proteasome system (UPS), including the only known OMM deubiquitinase Usp30^9,11^, and AAA ATPase ATAD1(ATPase family AAA domain containing 1)/Msp1 were also recently linked to peroxisome homeostasis^27,28^. Usp30 was originally reported to control mitochondrial fusion^10,29,30^, and Parkin-mediated mitophagy^29^.

Here we report that MARCH5 influences Pex26-dependent peroxisome formation. Pex26 is essential for forming the AAA ATPase complex (PEX1/PEX6) that facilitates protein import into peroxisomes. The data show that subcellular localization and Pex26 turnover are directly controlled by MARCH5 activity, providing further evidence on the direct role of the outer mitochondrial membrane in *de novo* peroxisome biogenesis. The data support the model that in peroxisome-containing cells, MARCH5 acts as a peroxisome biogenesis factor, while upon defective peroxisome biogenesis, as in Zellweger syndrome cells, it protects mitochondria from potentially toxic accumulation of peroxins on the OMM.

## Results and Discussion

### MARCH5 controls expression levels of Pex26 and PMP70 in peroxisome biogenesis-deficient cells

We and others reported that the predominantly outer mitochondrial membrane (OMM)-localized integral E3 Ub ligase MARCH5^3,7,8,22^ is required for *de novo* peroxisome biogenesis^3,4^. These works also supported the direct role of mitochondria in *de novo* peroxisome formation, initially reported by Sugiura *et al.*^2^. We set out to further test the importance of MARCH5 in peroxisomes. Global proteomic analyses of total cell lysates from WT and MARCH5^−/−^ cells revealed that “peroxisome protein import” is among the pathways significantly affected by MARCH5 knockout (Fig. 1A). Within this pathway, several classes of proteins were significantly downregulated, including proteins that mediate peroxisomal protein import through interaction with peroxisomal membranes (Pex1, Pex14, Pex5) and luminal proteins that are involved in regulation of peroxisomal lipid metabolism (ACAA1, ACOX1, AGPS, DECR2, and LONP2) among others (Fig. 1A, Fig S1A). Protein–protein interaction network analysis demonstrated a tightly connected module among these proteins, indicating coordinated impairment of proteins associated with peroxisome protein import pathway (Fig. S1B). Supporting the candidate protein assays (Fig. 1B-D and Ref^3^), the data show that steady-state expression of several peroxisome biogenesis factors (peroxins or Pex proteins) and other peroxisomal membrane proteins was reduced in MARCH5^−/−^ cells (Fig. 1A). Consistent with MARCH5’s role in peroxisome biogenesis, we also found that MARCH5 deficiency results in a significant reduction of luminal peroxisomal proteins expression levels (Fig. 1A). These data validate a critical role of MARCH5 in peroxisome biogenesis. In this work, we aimed to determine the mechanism and MARCH5 client proteins involved in this process.

**Figure 1:**
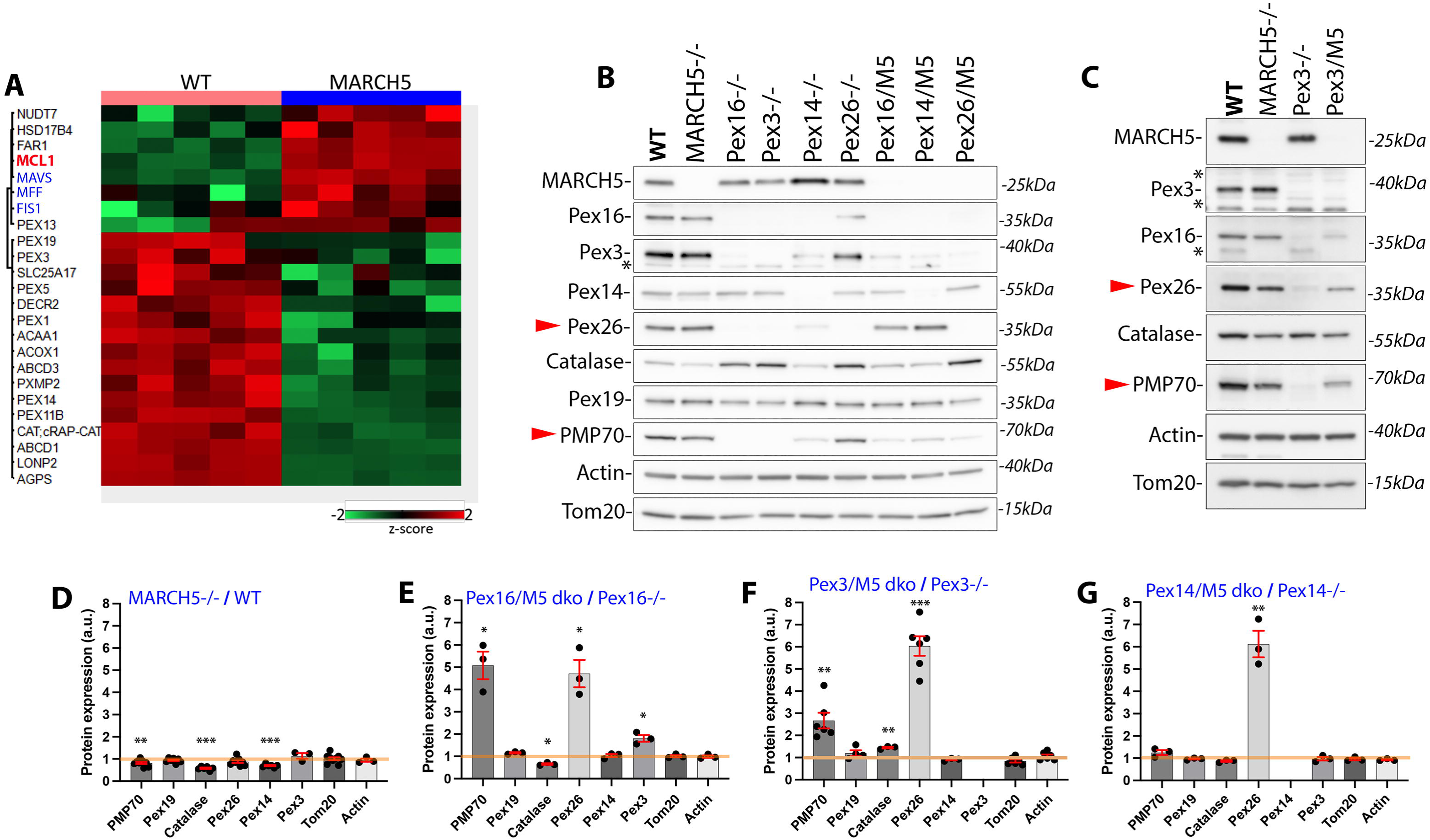
MARCH5 controls levels of Pex26 in peroxisome-biogenesis deficient cells. (**A**) Total cell lysates from WT and MARCH5^−/−^ HeLa cells (n = 5 biological replicates per group) were analyzed by mass spectrometry–based proteomics to evaluate steady-state protein expression changes in MARCH5^−/−^ compared to WT. Heatmap depicting differentially expressed proteins, where red denotes upregulation and green denotes downregulation. Mcl1 (red) is a MARCH5 client protein, MAVS, Mff, and Fis1 (blue) show mitochondrial and peroxisomal localizations. (**B**,**C**) Total cell lysates from the cells indicated in the figure were subjected to Western blot analysis to test the expression levels of representative peroxins. Actin and Tom20 are loading controls. **\***-non-specific or remaining after reblotting. Red arrowheads point to proteins markedly reduced in single Pex knockouts and protected by MARCH5 knockout in Pex/MARCH5 dko cells. (**D**-**G**) Relative protein expression levels in MARCH5-deficient *versus* MARCH5-expressing cells. (**D**) MARCH5^−/−^ *versus* WT, (**E**) Pex16/MARCH5 dko *versus* Pex16^−/−^, (**F**) Pex3/MARCH5 dko *versus* Pex3^−/−^, and (**G**) Pex14/MARCH5 dko *versus* Pex14^−/−^. The densitometric values obtained in MARCH5-deficient cells were divided by those obtained in MARCH5 expressing cells (e.g., MARCH5^−/−^ by WT cells). Mean ± SEM. N=3-9 independent experiments. Statistics (**D**-**G**): one-sample t-test and Wilcoxon signed-rank test versus 1 (value expected if there was no difference in protein expression). * p<0.05, **p<0.01, *** p<0.001.

Mutations or knockdowns of peroxisome biogenesis factors (peroxins or Pex-proteins), including Pex3, Pex16, and Pex14, lead to a significant reduction in expression of other peroxins^3,31,32^. To determine whether MARCH5 is critical for peroxin steady state expression levels in peroxisome-containing and peroxisome-deficient HeLa cells, we analyzed total cell lysates from WT (normal peroxisomes), MARCH5^−/−^ (increased number of Catalase-deficient peroxisomes^3^), Pex3^−/−^ (no detectable peroxisomes^2–4,8^), Pex16^−/−^ (no detectable peroxisomes^3^), and Pex14^−/−^ (accumulation of Tom20-positive, enlarged, mitochondria-derived pre-peroxisomes^3^), and respective cells also depleted of MARCH5 for steady state protein levels of candidate peroxins and other peroxisomal proteins (Fig. 1B-G). As we reported^3^ and as validated by the proteomic data shown in Fig. 1A, MARCH5 knockout leads to a reduction in the levels of several peroxisomal proteins, including catalase and Pex14, compared to WT cells (Fig. 1A,C).

Levels of Pex26 and peroxisomal fatty-acid transporter PMP70 were markedly reduced in the tested Pex-knockout cells (Fig. 1B,C). MARCH5 depletion in these peroxisome biogenesis-deficient cells led to significant stabilization of Pex26 (Fig. 1B-G), while PMP70 was stabilized in all tested cells except for Pex14/MARCH5 dko (Fig. 1B,G). Other peroxins, including Pex14 and Pex3, proteins earlier implicated in the generation of mitochondria-derived pre-peroxisomes^2,4^ and shown to localize to the mitochondria in peroxisome biogenesis deficient cells^2–4^, and Pex19, import receptor distributed between the cytoplasm and peroxisomes^33^, reported to recruit MARCH5 to peroxisomes in MARCH5-dependent pexophagy^8^, were affected to a lesser degree (Fig. 1B-G).

### Proteasomal degradation of Pex26 in peroxisome-deficient cells is controlled by MARCH5

We investigated the role of MARCH5 in regulating Pex26 protein levels. First, to test the degradation rates of Pex26, we applied the protein translation inhibitor cycloheximide (CHX). WT, MARCH5^−/−^, Pex3^−/−^, Pex14^−/−^, Pex16^−/−^ and respective Pex/MARCH5 dko cells were treated with CHX for 0 to 6hr, followed Western blot and densitometric quantification of tested proteins. We found that in contrast to WT and MARCH5^−/−^ cells, where Pex26 expression levels did not notably change within the 6hr time frame of the experiment (Figs. 2A,B and S2A,B), Pex26 was markedly reduced in Pex3^−/−^ (Fig. 2A,B), Pex16^−/−^ (Fig. S2A,B), and to lesser degree Pex14^−/−^(Fig. 2B). The less pronounced rates of Pex26 degradation detected in pre-peroxisome containing Pex14^−/−^ cells (Fig. 2B), as compared to peroxisome-deficient Pex3^−/−^ and Pex16^−/−^cells (Figs. 2A,B, and S2A,B, respectively), suggest that MARCH5 facilitates degradation of mitochondria-localized Pex26, and it is protected from degradation when it localizes, at least partially, on peroxisomes, as in Pex14^−/−^ cells^3^. Consistent with this notion, Pex26 has been reported to localize to the mitochondria when peroxisome biogenesis is deficient^15,18,28^. Indicating the critical role of MARCH5 in Pex26 turnover in peroxisome biogenesis-deficient cells, MARCH5 depletion completely inhibited Pex26 degradation in Pex3/MARCH5 dko, Pex16/MARCH5 dko, and Pex14/MARCH5 dko cells (Figs. 2A,B, and S2A,B). On the other hand, PMP70 degradation does not appear to be directly controlled by MARCH5 (Fig. 2A,D). The changes in PMP70 expression levels in Pex knockout cells can be indirectly linked to reduced Pex26 levels and are consistent with the fact that certain disease-linked mutations in Pex26 lead to decreased stability of this protein, affecting the overall peroxisomal structure and altering the normal PMP70/catalase ratio.^34^

**Figure 2:**
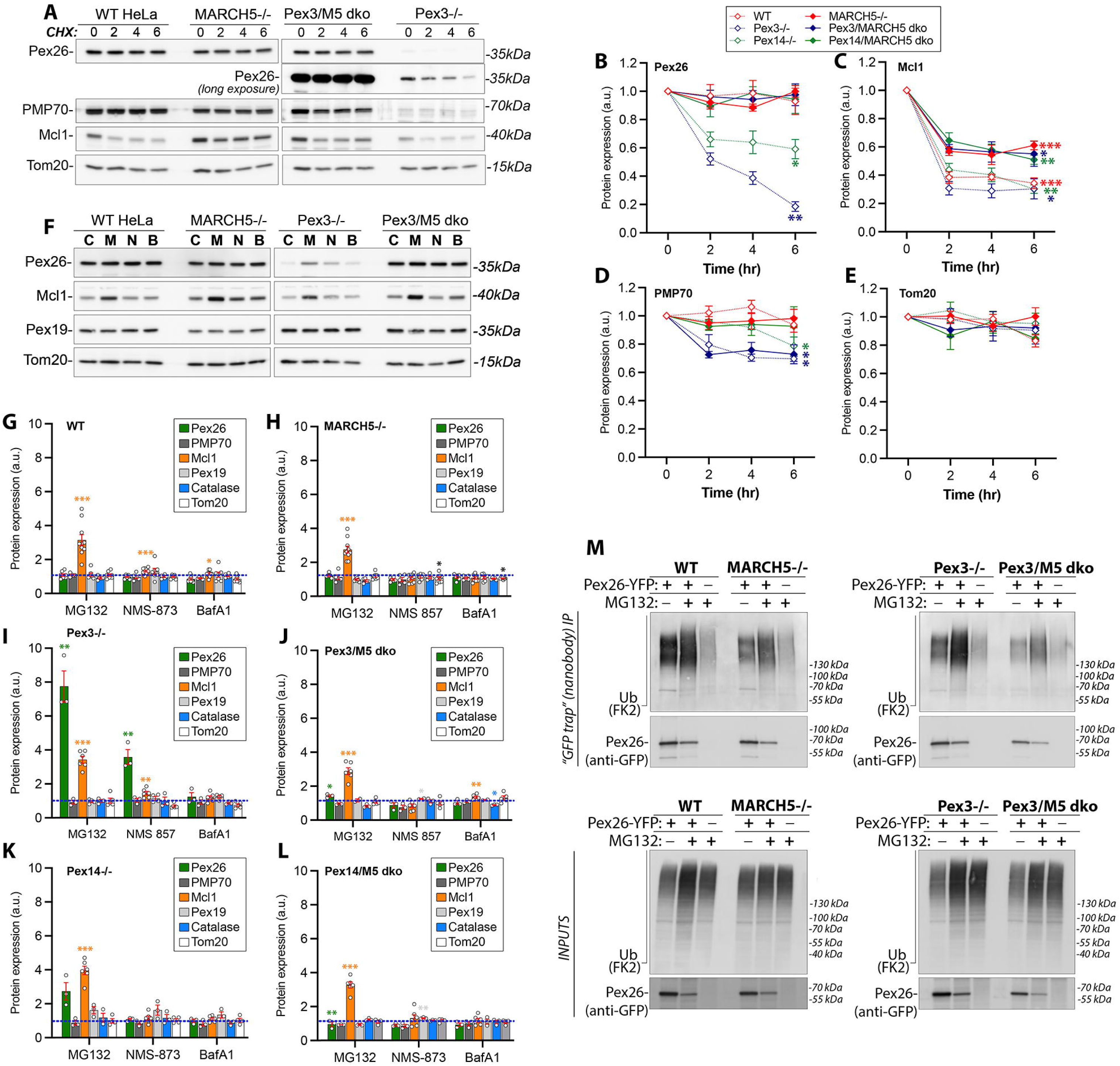
MARCH5 controls turnover rates of Pex26 in a proteasome and peroxisome-dependent manner. (**A**) Total cell lysates from WT, MARCH5^−/−^, Pex3^−/−^, and Pex3/MARCH5 dko HeLa cells treated with Cycloheximide (CHX; protein synthesis inhibitor) for 0 (Control), 2, 4, and 6 hr, and analyzed with Western blot as indicated in the figure. Tom20 is a loading control. (**B-E**) Quantifications of the expression of representative peroxisomal proteins in cells treated as in **A**. Expression levels of Pex26 (**B**), Mcl1 (**C**), PMP70 (**D**) and Tom20 (**E**) in cells indicated in the figure were quantified. Values obtained in vehicle (DMSO-treated cells; indicated as 0 hr) were taken as 1. Mean ± SEM. Statistics (**B**-**E**): paired t-tests comparing each timepoint to 0 h within condition, with Benjamini-Hochberg FDR correction. Two-way ANOVA (time/times condition): p(time)=4.23e-36, p(condition)=8.07e-12, p(interaction)=0.00304. Stars denote FDR-corrected significance (* p<0.05, ** p<0.01, *** p<0.001). **(F)** WT, MARCH5^−/−^, Pex3^−/−^ and Pex3/MARCH5 dko HeLa cells were treated with MG132 (proteasome inhibitor), NMS 873 (p97 inhibitor), and Bafilomycin A1 (BafA1; autophagy inhibitor) for 6 hr and analyzed with Western blot as indicated in the figure. Tom20 is a loading control. (**G**-**L**) Densitometric evaluation of the expression of representative peroxisomal proteins in cells treated as in **F** in WT (**G**), MARCH5^−/−^ (**H**), Pex3^−/−^ (**I**), Pex3/MARCH5 dko (**J**), Pex14^−/−^ (**K**), and Pex14/MARCH5 dko (**L**) HeLa cells. Values obtained in vehicle (DMSO; Control)-treated cells were taken as 1. Mean ± SEM. N=3-7. Statistics (**G**-**L**): paired t-tests comparing each drug treatment to control within the tested protein. * p<0.05, ** p<0.01, *** p<0.001. (**M**) Ubiquitination status of ectopically expressed Pex26-YFP in untreated and MG132-treated cells. WT, MARCH5^−/−^ (left panels), Pex3^−/−^, and Pex3/MARCH5 dko (right panels) HeLa cells were transfected with Pex26-YFP and treated with MG132 or vehicle (DMSO) for 5 hr and subjected to immunoprecipitation under denaturing conditions with GFP-trap nanobodies. Samples were analyzed for Ub with anti-conjugated Ub (FK2) antibody, Pex26 was detected with anti-GFP antibody. Controls are samples from MG132-treated untransfected cells. Protein levels in the respective inputs are shown in the bottom panels.

Next, we investigated the mechanism of Pex26 degradation. MARCH5 is an E3 Ub ligase with a role in proteasomal degradation of mitochondrial proteins, such as Mcl1 and MiD49^21,22,25^. MARCH5 has also been implicated in mitophagy and pexophagy^8,35^. We applied MG132, an inhibitor of the proteasome-dependent protein degradation, NMS-873^36^, an inhibitor of AAA-ATPase p97 (implicated in retrotranslocation and proteasomal degradation of mitochondrial membrane proteins^37,38^), and the autophagy inhibitor Bafilomycin A1 (BafA1). Cells were treated with the above-described compounds for 6hr followed by Western blot analysis (Figs. 2F-L and S2C). Pex26 protein levels were markedly increased in MG132-treated Pex3^−/−^(Fig. 2F,I), Pex16^−/−^ (Fig. S2C), and to a lesser degree Pex14^−/−^ cells (Fig. 2K), as compared to MG132-treated WT, MARCH5-/-, and Pex/MARCH5 dko cells (Figs. 2G,H,J,L, and S2C), where no Pex26 increases were detected. Thus, it is likely that peroxisome localization of Pex26 (as in WT and MARCH5^−/−^ cells) or lack of MARCH5 (as in MARCH5^−/−^ and Pex/MARCH5 dko cells) reduces Pex26 dependence on proteasomal degradation. Furthermore, p97 inhibition increased Pex26 levels only in Pex3/M5 dko cells, but not in Pex14/MARCH5 dko, WT, and MARCH5^−/−^ cells, indicating that p97 participates in Pex26 degradation in peroxisome-deficient cells, but its role is less pronounced. It is possible that ATAD1, another AAA-ATPase implicated in the removal of peroxisomal proteins from the OMM^15,18^, also contributes to the retrotranslocation step and thereby rescues p97 deficiency.

Notably, Mcl1, a MARCH5 client protein^21,25,26^, was stabilized by MG132 in all tested cells (Fig. 2F-L), and its turnover was not affected by peroxisome status (Fig. 2A,C), indicating that differences in Pex26 expression are not due to aberrant MARCH5 activity or dysfunction of the OMM-localized protein degradation machinery. Since inhibition of autophagy with BafA1 did not show any effects (Fig. 2G-L), one can conclude that proteasomal degradation of Pex26 is a major mechanism regulating its expression levels. Consistent with this notion, analyses of the ubiquitination status of Pex26 in WT, MARCH5^−/−^, Pex3^−/−^, and Pex3/M5 dko cells show higher Ub signal in MARCH5-expressing cells (Fig. 2M). However, despite low turnover rates of Pex26 in WT cells, high ubiquitination levels of this protein were apparent (Fig. 2M), suggesting that a function other than the proteasomal degradation of Pex26 is controlled by MARCH5 E3 Ub ligase activity.

To sum up, in contrast to WT and MARCH5^−/−^ cells in which Pex26 is stable, turnover rates of Pex26, including synthesis and degradation, are increased in peroxisomal biogenesis-deficient Pex3^−/−^, Pex16^−/−^, and Pex14^−/−^ cells. Pex26 degradation in these cells is controlled by MARCH5. High-turnover rate proteins, such as anti-apoptotic Bcl2 family protein Mcl1 and transcription factors, are critical for rapid cellular adaptation. The increases in the turnover rates of Pex26 in peroxisome biogenesis-deficient cells (Figs. 2A-L, S1A-C) suggest that Pex26 potentially converts into a signaling protein in the absence of peroxisomes, and only when these organelles are present, Pex26 can assemble in correct peroxisomal complexes and lose its unstable nature. Consistent with the potential Pex26 signaling role, an increase in Pex26 expression levels in WT and MARCH5^−/−^ cells treated with Torin1, a potent and selective ATP-competitive inhibitor of mTORC1 and mTORC2 activity^39^, was detected, while other tested proteins were not affected or reduced (Fig. S1H-J). Torin1 induces macroautophagy^40^, including pexophagy^8,41^, accelerates proteasomal degradation of long-lived proteins^40^, and reduces translation^42^. The accumulation of Pex26 in Torin1-treated cells may indicate protection from mTOR inhibition-dependent degradation and a potential role of this protein in controlling peroxisome biogenesis signaling. This interesting possibility will be tested in the subsequent studies.

### The interaction of Pex26 and MARCH5 influences the subcellular localization of Pex26

Immunoprecipitation of MYC-tagged WT MARCH5 (MYC-WT M5), RING domain mutant MARCH5 (MYC-M5H43W), and C-terminal truncation MARCH5 mutant (MYC-M5deltaC) tested interactions of MARCH5 with Pex26, and other candidate peroxins (Fig. 3A; not shown). The latter two constructs were shown to eliminate MARCH5-dependent proteasomal degradation of its client proteins Mcl1 and MiD49^21,22^. All tested MYC-MARCH5 proteins coimmunoprecipitated endogenous Pex26 (Fig. 3A) in MARCH5^−/−^ cells, while, likely due to low expression, no interaction of Pex26 with tested MYC-MARCH5 proteins was detected in Pex3/MARCH5^−/−^cells. In contrast mitochondrial fission factor (Mff), a protein we reported to bind MARCH5 in HCT116 cells^21^, interacted with MARCH5 in both tested cell lines (Fig.3A). Thus, despite high stability of Pex26 in WT and MARCH5^−/−^ cells (Fig. 2A-D), a robust interaction of Pex26 with active and inactive MARCH5 variants, suggest that other than proteasomal degradation of Pex26 MARCH5-dependent activity occurs in peroxisome-containing cells. No clear interaction between MYC-MARCH5 proteins and Pex14 and Pex19 was detected, despite high expression levels of these proteins and high-quality antibodies (not shown).

**Figure 3:**
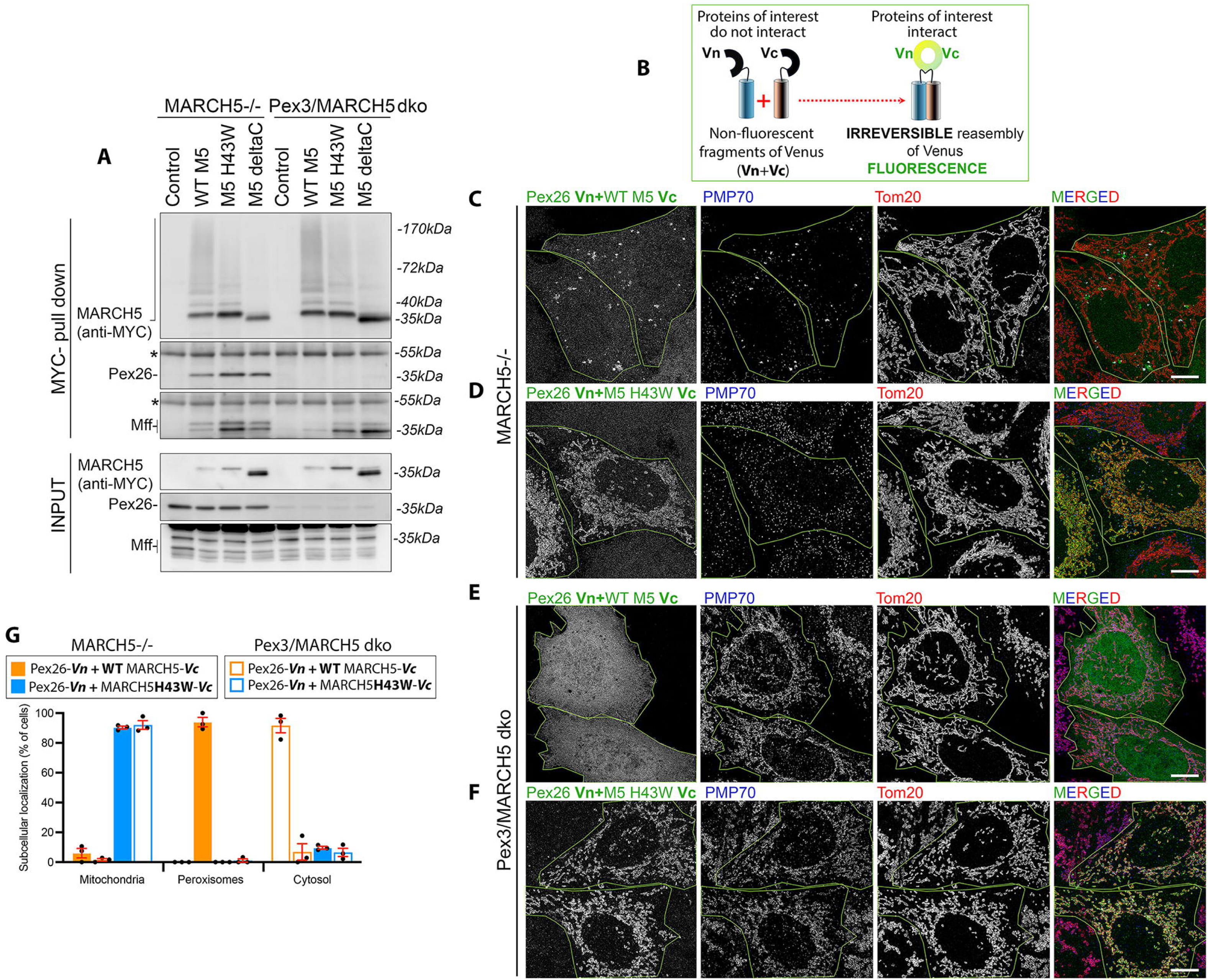
MARCH5-dependent control of Pex26. (A) Total cell lysates obtained from control-, MYC-MARCH5–, MYC-MARCH5^H43W^–, and MYC-MARCH5d^eltaC^–transfected MARCH5^−/−^ and Pex3/MARCH5 dko HeLa cells were subjected to MYC immunoprecipitation. Immunoprecipitated samples (top panels) and inputs (bottom panels) were analyzed by Western blot as indicated. (B) Overview of bimolecular fluorescence complementation (BiFC). Split N-terminal (Vn) and C-terminal (Vc) fragments of fluorescent protein Venus fused to proteins of interest. (**C**-**F**) Typical images of MARCH5^−/−^ (**C**,**D**), and Pex3/MARCH5 dko (**E**,**F**) HeLa cells expressing Pex26-Vn and WT MARCH5-Vc (**C**,**E**), or Pex26-Vn and MARCH5^H43W^-Vc (**D**,**F**), fixed at 16-18 hr after transfections. The BiFC signal is green in the merged images. Cells were also immunostained to detect peroxisomes (PMP70; blue) and mitochondria (Tom20; red). Bars:10μm. Cells showing the BiFC signal are overlaid with green lines. (H) Quantification of subcellular localization of BiFC signal in MARCH5^−/−^ and Pex3/MARCH5 dko cells. N=3 independent experiments (each time >80 cells were counted per condition). Mean ± SEM.

Next, we tested the subcellular location of Pex26 bound to MARCH5 (Fig. 3B-G). Split fluorescent proteins, such as GFP or yellow fluorescent protein Venus (used in this study), have a very high binding affinity, with dissociation constants in the sub-nanomolar to low nanomolar range^43,44^. The strong interaction of split N-terminal and C-terminal fragments allows for the reassembly of the fluorescent protein (bimolecular fluorescence complementation; BiFC), making it ideal for monitoring localization of protein-protein interactions within cells with high spatial resolution^45,46^. The stable nature of reassembled Venus fragments would also stabilize transient complexes of tested proteins and perhaps reveal the intermediate steps in the analyzed pathway (Fig. 3B). We reasoned that BiFC should reveal interactions between MARCH5 and peroxisome proteins, if they occur, and their subcellular location. First, we tested the BiFC signal in MARCH5^−/−^ (Fig. 3C,D), and Pex3/MARCH5 dko (Fig. 3E,F) cells expressing N-terminal Venus fragment (Vn) fused to Pex26 (Pex26-Vn), together with C-terminal Venus fused to WT MARCH5 (WT M5-Vc; Fig. 3C,E), or M5 H43W-Vc (Fig. 3D,F). In MARCH5^−/−^ and Pex3/MARCH5 dko cells co-expressing Pex26-Vn and MARCH5^H43W^-Vc, BIFC signal was predominantly mitochondrial (Figs. 3D,F,G and S3F,H). On the other hand, Pex26-Vn and WT MARCH5-Vc combination showed BiFC signal colocalizing with peroxisome markers PMP70 and Pex14 (Figs. 3C,G, and S3E,G, respectively). Thus, newly expressed Pex26 is targeted to the peroxisomes only in the presence of WT MARCH5. Accumulation of the Pex26-Vn/MARCH5H43W-Vc complexes on the OMM indicates that MARCH5 E3 Ub ligase activity is vital for the transfer of Pex26 to peroxisomes. The cytosolic BiFC signal of Pex26-Vn + WT MARCH5-Vc detected in peroxisome-deficient Pex3/MARCH5 dko cells (Fig. 3E,G) indicates either cytosolic binding of these proteins or redistribution of reformed Venus to the cytosol due to WT MARCH5-Vc-dependent degradation of Pex26-Vn. Considering that in Pex3-deficient cells MARCH5 controls Pex26 degradation (Fig. 2A,B,I), the latter possibility is more likely. E3 Ub ligases, including MARCH5, are known for the “safety valve” mechanism in which their oligomerization-induced self-degradation^47,48^ protects cells from excessive degradation of their target proteins. Like Pex26-Vn + WT MARCH5-Vc, WT MARCH5-Vn + WT MARCH5-Vc BiFC signal was detected in the cytosol (Fig. S3A,C), suggesting cytosolic localization of degradation products. Importantly, as we showed with MYC-tagged MARCH5 constructs^21^, dramatically reduced levels of MARCH5’s client protein Mcl1^21,25,26^ were apparent in Pex26-Vn + WT MARCH5-Vc, but not Pex26 Vn + MARCH5^H43W^-Vc expressing cells (Fig. S3G-I), indicating that MARCH5 activity is not impeded by stable interaction with Pex26.

### MARCH5 controls the subcellular localization of newly expressed peroxins

The data (Fig. 3) suggest that MARCH5 is vital for the transfer of newly synthesized Pex26 to peroxisomes. The accumulation of Pex26 on the mitochondria in MARCH5^H43W^-expressing cells indicates that Pex26 transfers through the OMM on its way to peroxisomes in a MARCH5-dependent manner. It is well established that under experimental or disease-associated inhibition of peroxisomal biogenesis, some tail-anchored peroxins, including Pex14 and Pex26, accumulate on the OMM^3,17,19,49,50^. It has been proposed that these proteins mislocalize to the OMM due to the absence of peroxisomes^17,19^. However, we found that a multi-TM domain protein, PMP70, also localizes to mitochondria in peroxisome-deficient cells (Fig. 3F and Ref^3^), suggesting that not only tail-anchored single TM domain peroxins, but also other integral peroxisomal proteins can be found on the mitochondria.

We further investigated whether mitochondria serve as an intermediate location in the delivery of newly generated peroxins to peroxisomes in peroxisome-containing cells. We reasoned that if a newly generated peroxin is primarily detected on peroxisomes in WT cells but localizes to the mitochondria in MARCH5^−/−^ cells, its delivery to peroxisomes likely requires the activity of MARCH5, supporting the role of MARCH5 and mitochondria in this pathway. Other localizations (e.g., ER or cytosol) would indicate participation of the non-mitochondrial component of *de novo* peroxisome biogenesis (Fig. 4A). While ectopic expression of fluorescent protein-fused protein of interest can lead to “overexpression” artifacts, this approach is ideal for testing the role of MARCH5 in the transfer of newly synthesized peroxins to peroxisomes. Overexpressed peroxins (even at the lowest expression levels, as we analyzed) are likely to overcome their transport system(s) and reveal the intermediate locations on their way to peroxisomes.

**Figure 4:**
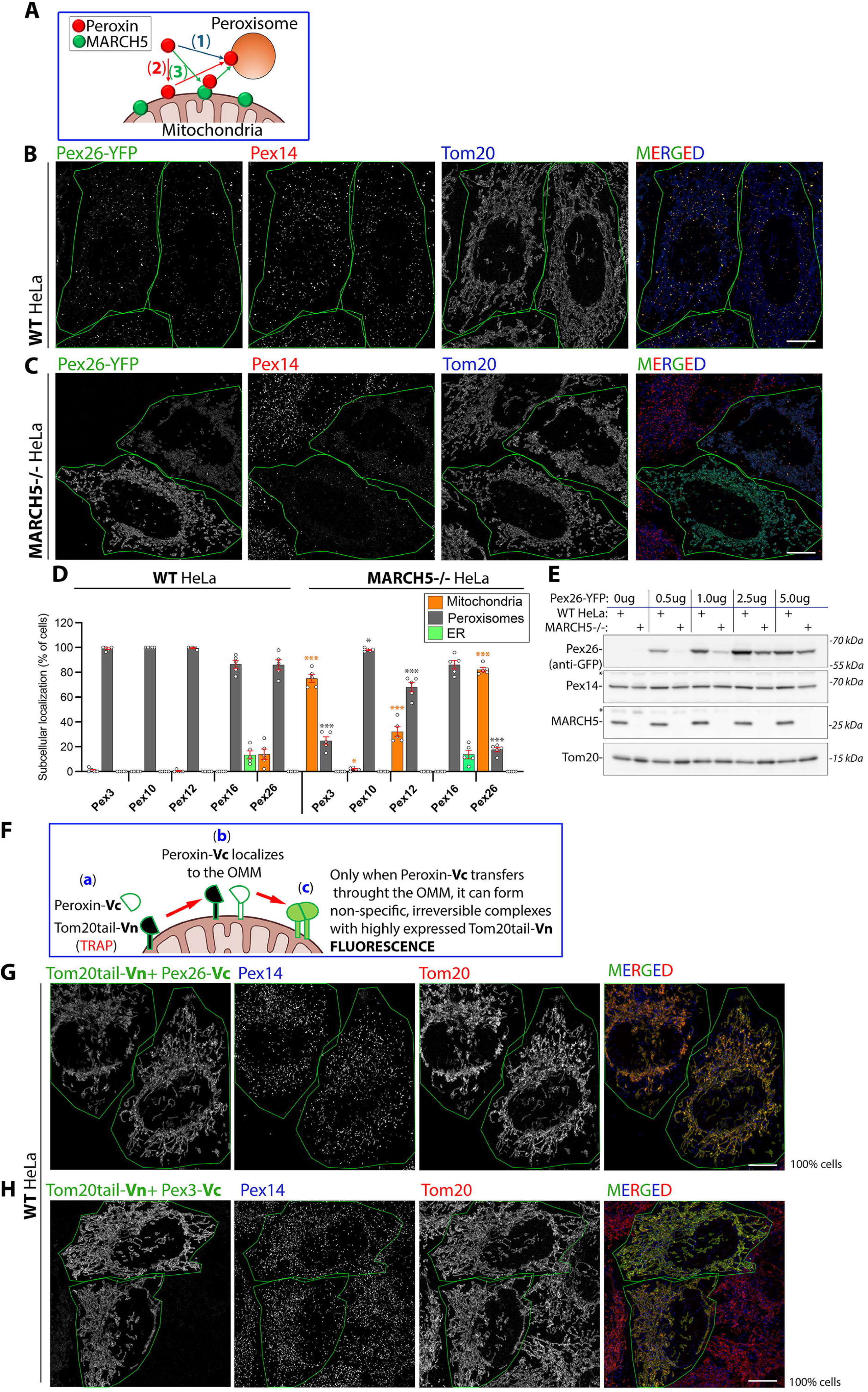
MARCH5 controls the subcellular localization of newly expressed peroxins. (**A**) Newly synthesized peroxins could be delivered to peroxisomes through (**1**) mitochondria-independent pathway (direct or through ER; no mitochondrial accumulation of the candidate protein detected under any experimental conditions), (**2**) mitochondria but without contribution of MARCH5 (accumulation of candidate proteins detected in peroxisome-deficient cells, but not affected by MARCH5 activity), and (**3**) MARCH5-dependent pathway (accumulate on mitochondria and affected by MARCH5 activity). (**B**, **C**) Images of Pex26-YFP (green)-expressing WT (**B**) and MARCH5-/- (**C**) HeLa cells analyzed at 14 hr after transfection. Cells were immunostained to detect peroxisomes (Pex14, red) and the OMM (Tom20, blue). Bars: 10 μm. Pex26-YFP-expressing cells are overlayed with green lines. **(D)** Quantification of subcellular localization of YFP-tagged Pex3, Pex10, Pex12, Pex16, and Pex26 imaged as in **B**,**C**. Mean ± SEM. N=5-7. Statistics (**D**): Paired t-tests comparing each localization in MARCH5^−/−^ cells to WT HeLa cells, with Benjamini-Hochberg FDR correction. FDR-corrected significance (* p<0.05, ** p<0.01, *** p<0.001). **(E)** Western blot analysis of expression levels of Pex26-YFP in WT and MARCH5^−/−^ HeLa cells. Four different amounts of Pex26-YFP plasmid (0.5 μg, 1.0 μg, 2.5 μg, and 5 μg per 10 cm tissue culture plate) were transfected into WT and MARCH5^−/−^ cells and analyzed 14 hr after transfection. Tom20 is a loading control. * non-specific. **(F)** Peroxins fused with a split yellow fluorescent protein Venus fragment Vc (Peroxin-Vc) (**a**) would, though the interaction of split Venus fragment Vc, with Vn fused to highly expressed OMM-localized Tom20tail (Tom20tail-Vn; **a**), form non-specific, irreversible complexes (**b**), and develop BiFC fluorescence (**c**), only if they transfer through the OMM. (**G, H**) WT HeLa cells transfected with BiFC constructs, Tom20tail-Vn and Pex26-Vc (**G**), Tom20tail-Vn and Pex3-Vc (**H**) (cells showing BiFC signal are overlaid with green lines; BiFC is green on merged images), were immunostained to detect peroxisomes (Pex14; blue), and the OMM (Tom20; red). Bars: 10 μm. All 100% cells showed mitochondrial localization of the BiFC signal in **G** and **H**. N = 3 (>80 cells counted each time).

WT and MARCH5^−/−^ HeLa cells were transfected with yellow fluorescent protein (YFP)-tagged peroxins, Pex3-YFP, Pex10-YFP, Pex12-YFP, Pex16-YFP, and Pex26-YFP, and analyzed for their subcellular localization at ∼14 hours after transfection (Fig. 4B-D). In WT HeLa cells, Pex3-YFP, Pex10-YFP, and Pex12-YFP predominantly localized to peroxisomes, while Pex26-YFP showed major distribution to peroxisomes and some mitochondrial localization (Fig. 4B-D). On the other hand, in MARCH5^−/−^ cells, Pex3-YFP, and Pex26-YFP were found predominantly on the mitochondria, while mitochondrial localization of Pex12-YFP was also significantly increased (Fig. 4D). The reported ER localization of GFP-tagged Pex16^2,51^, was only detected when it was highly overexpressed in both WT and MARCH5^−/−^cells, while it invariably showed peroxisomal localization in cells expressing moderate or low levels of this protein (Fig. 4D). Other tested peroxins did not show any ER localization, regardless of expression levels. Importantly, levels of expressed Pex26-YFP were comparable in WT and MARCH5^−/−^ cells, regardless of the amounts of transfected plasmids (Fig. 4E). We tested a range of 0.5 to 5μg of Pex26-YFP plasmid DNA transfected in standard 10-cm tissue culture plates; with ∼2.5 μg per plate being equivalent to imaging transfections.

Since BiFC leads to irreversible stabilization of protein complexes^41,4^ at the sites they are formed, we reasoned that a combination of high levels of Vn- or Vc-fused OMM marker and Vn-or Vc-peroxin should result in a non-specific BiFC signal on the OMM if the given peroxins transfer through the mitochondria. In this assay, any OMM-interacting protein would generate a BiFC signal with a highly expressed integral OMM protein (Venus molecular trap; Fig. 4F). Indeed, we found that in WT HeLa cells co-expressing 30 N-terminal amino acid residues of the OMM protein Tom20 fused with Vn (Tom20-tail-Vn) with Pex-26-Vc, or Pex3-Vc (ratio of plasmids used in transfection 3:1), the BiFC signal was restricted to the OMM in 100% of the cells (Fig. 4G,H). Since the combination of Tom20-tail-Vn and Tom20-tail-Vc also resulted in a highly restricted OMM BiFC signal, but single expression of these constructs did not generate detectable fluorescence (not shown), the specificity of this assay is high.

These observations indicate that some peroxins (e.g., Pex3 and Pex26) transfer through mitochondria to arrive at their destination at peroxisomes even when mature peroxisomes are present, as in WT and MARCH5^−/−^ cells rescued with WT MARCH5, providing evidence countering the assumption that peroxisomal proteins localize to the mitochondria just due to the absence of their target organelles^17,19^.

### MARCH5 controls Pex26-dependent peroxisome biogenesis

To test the degree to which MARCH5 controls Pex26-dependent peroxisome biogenesis, we investigated peroxisomal status in CRISPR/Cas9-generated Pex26^−/−^ and Pex26/MARCH5 dko HeLa cells (Fig. 5, see Figs. 1B and 5K for Western blot validation). The mosaic peroxisome morphology was detected in Pex26^−/−^ and Pex26/MARCH5 dko cells (Fig. 5A-G), with ∼50% of the cells showing normal peroxisomes, judged by peroxisomal localization of luminal protein Catalase (Fig. 5G) and peroxisome number (Fig. 5I), while ∼50% of the cells showed cytosolic localization of Catalase (Fig. 5G). Similar phenotypic mosaicism, with different clones of peroxin-deficient cells varying in the number of cells containing peroxisomes, ranging from very low to high, was reported for Pex16^−/−^ HeLa cells^52^. The mechanism of the mosaic phenotypes in Pex26-and Pex16-deficient cells needs to be investigated.

**Figure 5:**
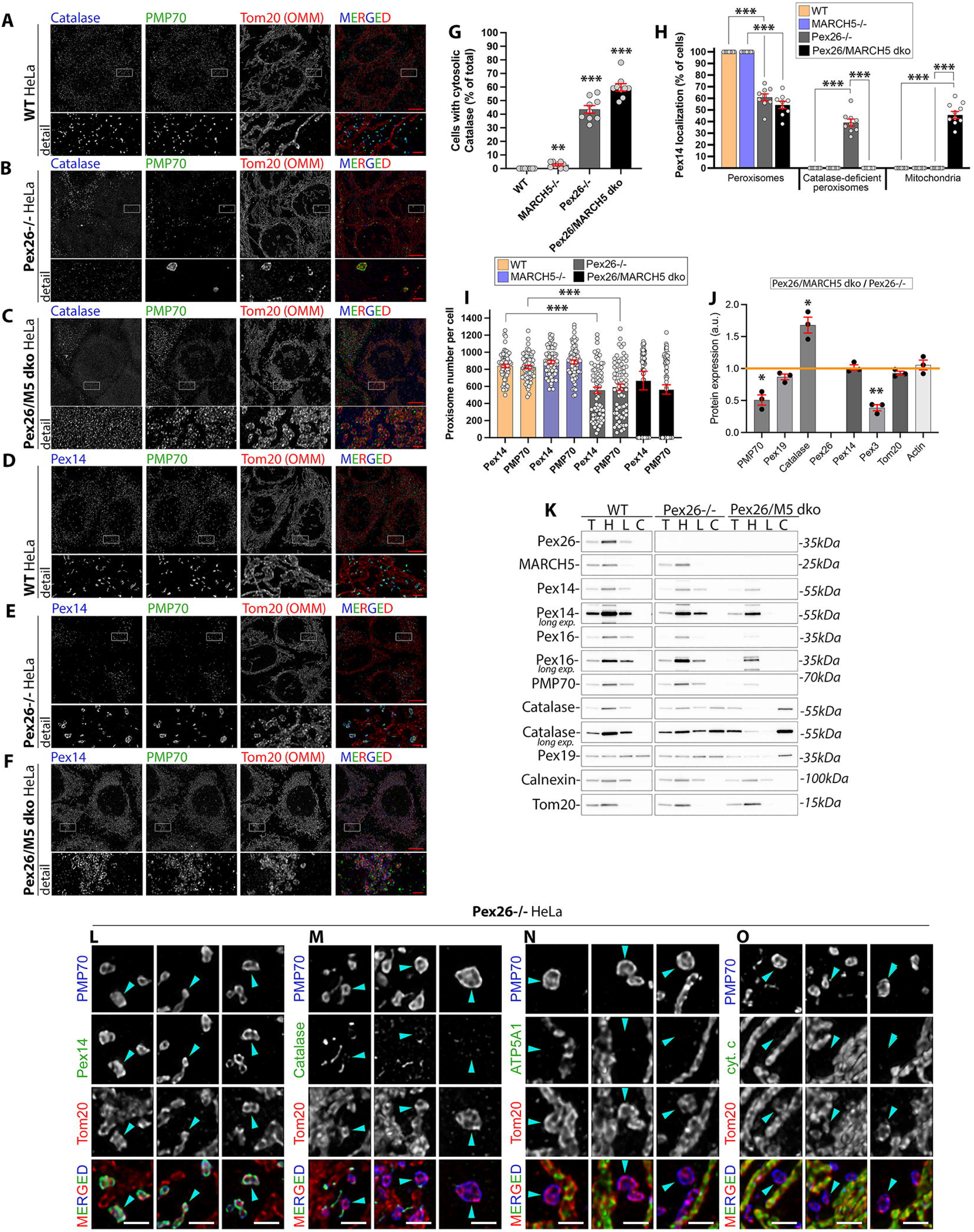
MARCH5 controls the formation of mitochondria-derived pre-peroxisomes in Pex26 knockout cells. (**A**-**F**) WT (**A**,**D**), Pex26^−/−^ (**B**,**E**), and Pex26/MARCH5 dko (**C**,**F**) HeLa cells were immunostained to detect catalase (**A**-**C**; blue), Pex14 (**D**-**F**; blue), PMP70 (**A**-**F**; green), and the OMM protein Tom20 (**A**-**F**; red). Bars: 10 μm and 2.5 μm (detail images). “Detail” images show areas marked with white rectangles. **(G)** Cytosolic localization of catalase in Pex26^−/−^ and Pex26/MARCH5 dko was detected by immunofluorescence. Cells were stained, as in **A**-**C**, and quantified in a blinded manner. Mean ± SEM. N=3 (100 cells counted in each independent experiment). Cells with cytosolic catalase were quantified as a percentage of total cells across four genotypes. Each genotype was compared to WT using unpaired t-tests and Wilcoxon rank-sum tests. Multiple comparisons were corrected using the Benjamini-Hochberg FDR method. Raw datapoints were displayed alongside mean ± SEM. **(H)** Subcellular localization of Pex14 in cells indicated in the figure was detected by immunofluorescence (as in **D**-**F**) and quantified in a blinded manner. Mean ± SEM. N=3 (n=9; 100 cells counted in 3 independent chambers in 3 independent experiments). Localization of Pex14 was quantified across subcellular compartments. Planned comparisons between genotypes were performed using unpaired Welch t-tests and Wilcoxon rank-sum tests. Multiple comparisons were corrected using the Benjamini-Hochberg FDR method. **(I)** Number of Pex14- and PMP70-positive peroxisomes/cell in cells indicated in the figure. Mean ± SEM. Data pulled from three independent experiments (n=60-70 cells/condition). Peroxisome number per cell was quantified using PEX14 or PMP70 as markers. Each point represents one cell. Comparisons between WT and knockout genotypes were performed using unpaired Welch t-tests and Wilcoxon rank-sum tests. Multiple comparisons were corrected using the Benjamini-Hochberg FDR method. **(J)** Relative protein expression levels in total cell lysates from Pex26/MARCH5 dko *versus* Pex26^−/−^ HeLa cells (see Fig. 2 for typical Western blot data). Mean ± SEM. N=3. Statistics (**G**-**J**): one-sample t-test and Wilcoxon signed-rank test versus Control (**G**-**I**), and *versus* 1 (**J**; value expected if there was no difference in protein expression). (* p<0.05, ** p<0.01, *** p<0.001). **(K)** WT, Pex26^−/−^, and Pex26/MARCH5 dko HeLa cells were fractionated into heavy-membrane (H), light membrane (L), and cytosolic (C) fractions, followed by a Western blot to detect proteins indicated in the figure. TCL, total cell lysates. Calnexin (ER membrane protein) and Tom20 (OMM integral protein) are loading controls. (**L**-**O**) Detail images from Pex26^−/−^ HeLa cells immunostained to detect peroxisomal membrane proteins PMP70 (**L**-**O**; blue) and Pex14 (**L**; green), peroxisomal luminal protein catalase (marker of mature peroxisomes, **M**; green), marker of mitochondrial inner membrane, ATP5A1 (**N**, green), mitochondrial intermembrane space marker, cytochrome c (**O**; green) and the OMM integral protein Tom20 (**L**-**O**; red). Arrowheads point to PMP70-positive structures and corresponding signals of other tested proteins. Bars: 1 μm.

Peroxisomes in Pex26^−/−^ cells showing cytosolic Catalase were enlarged and less numerous than in WT HeLa cells (Fig 5B,E,I). These enlarged, Catalase-deficient peroxisomes colocalized with the OMM marker Tom20 (Fig. 5K,N), but were deficient in inner mitochondrial membrane (IMM, ATP5A1; Fig. 5M), and intermembrane space (IMS; cytochrome c; Fig. 5N) markers, indicating that these structures are not fragmented mitochondria, but rather the OMM marker-containing mitochondria-derived pre-peroxisomes, similar to those we reported in Pex14*^−/−^* cells^3^. On the other hand, in Pex26/MARCH5 dko cells, cytosolic Catalase was invariably associated with the absence of detectable peroxisomes and with the mitochondrial localization of the tested peroxisome membrane proteins, PMP70 and Pex14 (Fig. 5C,F,H). Thus, it is likely that MARCH5 controls peroxisome formation upstream of Pex26, and underdeveloped peroxisomes accumulate due to MARCH5 activity, but in the absence of Pex26-mediated biogenesis steps.

Cell fractionation (Fig. 5K) verified the imaging data. Specifically, MARCH5 deficiency in Pex26^−/−^ cells led to marked or complete reduction of peroxins, including Pex14 and Pex16, and PMP70 in the light membrane fractions. Furthermore, reduction of the Catalase signal in heavy and light membrane fractions, and concurrent increase in cytosol, was also apparent in Pex26/MARCH5 dko cells (Fig. 5K). Validation of this assay for determination of peroxisome status is described in Ref ^3^.

### Concluding remarks

Recent evidence indicates that *de novo* peroxisome biogenesis requires mitochondria and depends on the activity of E3 Ub ligase MARCH5^2–4,53^. However, until now, the mechanism and mitochondrial factors implicated in the formation of mitochondrial subsets of peroxisomes have been mostly determined using peroxisome-deficient models. We report for the first time that mitochondria and in particular MARCH5 are vital for the *de novo* peroxisome biogenesis in peroxisome-containing cells. We also identified Pex26 as the MARCH5 client protein that in peroxisome-containing cells transiently associates with the OMM and is transferred to existing peroxisomes in a MARCH5-dependent manner. In peroxisome-deficient cells, however, MARCH5 targets Pex26 for proteasomal degradation in the cytosol. Thus, depending on peroxisomal status, MARCH5 acts as a peroxisome biogenesis factor or protects mitochondria from abnormal protein load on the OMM when peroxisome biogenesis is defective, and peroxins and other peroxisome membrane proteins localize to the OMM.

## Material and Methods

### Cells and cell culture

HeLa cells were maintained in Dulbecco Modified Eagle Medium (DMEM; Invitrogen) supplemented with 10% fetal bovine serum (FBS; Sigma), non-essential amino acids (Sigma), sodium pyruvate (Invitrogen), and penicillin/streptomycin (Invitrogen). Cells were maintained in 5% CO_2_ at 37°C, as reported ^3,21,54^.

### Knockout cells, DNA constructs, and transfections

The following cells lines were reported and validated earlier: MARCH5^−/−^ HeLa^3^, Pex3^−/−^ and Pex3/MARCH5 dko HeLa^3^, Pex16^−/−^ and Pex16/MARCH5 dko^3^, Pex26^−/−^, and Pex26/MARCH5 dko^3^. Mammalian expression vectors of MYC-tagged WT MARCH5, MARCH5^H43W^, and MARCH5^deltaC^ were reported^21,54^. Pex3-YFP was reported^3^. Pex16-GFP was provided by Dr. Haidi McBride (McGill University, Canada)^2^. Pex10-YFP, Pex12-YFP and Pex26-YFP were provided by Dr. Min Zhuang (Shanghai Institute of Technology)^4,8^. Mammalian expression vectors used in bimolecular fluorescence complementation (BiFC) assay were generated as follows. Plasmids pcDNA3-UiFC-N (Vn) and pcDNA3-UiFC-C (Vc)^45^ were cut with *Not*I-HF and *Cla*I to allow for insertion of new DNA fragments. WT MARCH5 was amplified from YFP-N1-MARCH5 vector^7^ using F1 (5’-AGATGAGCGGCCGCATGCCGGACCAAGCCCTACA-3’) and R1 (5’-TCATCTATCGATTGCTTCTTCTTGTTCTGGATAATTCAGAATTTTGC-3’). Plasmid YFP-N1-MARCH5^H43W 7^ was used as a template to generate MARCH5^H43W^ gene fragment. For amplification of Pex26 gene, an ORF clone (Biocat, #36414012-ABM) was used as template with oligonucleotides F2 (5‘-AGATGA GCGGCCGC ATGAAGAGCGATTCTTCGACCTCTG-3’) and R3 (5‘-TCATCTATCGATGTCACGGATGCGGAGCTGGTA-3’). Pex3 gene was amplified from Pex3-YFP with oligos F3 (5’-AGATGA GCGGCCGC ATGCTGAGGTCTGTATGGAATTTTCTGAAAC-3’) and R4 (5’-TCATCTATCGATCTTCTCCAGTTGCTGAGGGGTACTAAAA-3’All PCR products were cut *Not*I-HF, *Cla*I before ligation into Vn or Vc vectors. For addition of N-terminal Tom20 tail, oligonucleotides F4 (5’-GGCCGCATGGTGGGTCGGAACAGCGCCATCGCCGCCGGTGTATGCGGGGCCCTTTTCATTGGGTACTGCATCTACTTCGACCGCAAAAGACGAAGTAT-3’) and R5 (5’-CGATACTTCGTCTTTTGCGGTCGAAGTAGATGCAGTACCCAATGAAAAGGGCCCCGCATACACCGGCGGCGATGGCGCTGTTCCGACCCACCATGC-3’) were annealed, phosphorylated, and directly inserted into *Not*I-HF/*Cla*I-digested Vn and Vc vectors. All plasmid sequences were verified by Sanger sequencing (Microsynth).

For imaging experiments, cells were transfected with 0.2 μg DNA per well in 4 well Nunc Lab-Tek II Chambered Coverglass (culture area 1.7 cm^2^; Invitrogen), resulting in low expression of proteins of interest. Cells were transfected with Lipofectamine 3000 (Thermo Fisher Scientific) according to the manufacturer’s instructions. Cells were used at ∼14-20 h after transfection.

### Cell lysates, cell fractionation, and Western blot

For total-cell lysates, cells were collected by scraping into ice-cold PBS, washed, and suspended in ice-cold PBS. Cell suspensions (100-200 μl) were lysed in the same volumes of 2 x SDS sample buffer (Thermo Fisher Scientific) supplemented with 5% β-mercaptoethanol (Millipore) and incubated at 100°C for 10 min, as described^3,21,22^. Cell fractionation was performed as previously described^3,21,54^. Cells were washed once with ice-cold PBS and scraped into 15-ml tubes in ice-cold PBS; this was followed by centrifugation at 500 × g for 5 min. The cell pellets were resuspended in ∼3 volumes of fractionation buffer (10 mM HEPES, 10 mM NaCl, 1.5 mM MgCl_2_, 5 mM N-ethylmaleimide (NEM), and protease inhibitors (Sigma). Cells were then passed 15 times through a 25-G needle attached to a 1-ml syringe to disrupt cell membranes. This suspension was centrifuged at 2500 × g at 4°C for 5 min to remove unbroken cells and cell debris. The supernatant was centrifuged at 6000 × g at 4°C for 10 min to pellet the heavy membrane (HM) fraction. The supernatants were centrifuged at 21,000 × g at 4°C for 10 min to pellet the light membrane (LM) fraction. To reduce cytosolic and HM contaminations, the HM and LM fractions were washed in ice-cold PBS supplemented with protease inhibitors (Sigma) and recentrifuged at 8000 × g, and 21,000 × g, respectively. Protein concentrations were measured directly in the samples using a NanoDrop 1000 spectrophotometer (Thermo Fisher Scientific). 50 μg of protein per sample was separated on 4-20% gradient Novex Tris-glycine polyacrylamide gels (Thermo Fisher Scientific) and transferred onto polyvinylidene fluoride membranes (Bio-Rad Laboratories). Membranes were blocked in 5% blocking-grade nonfat dry milk (Bio-Rad Laboratories) or, for Ub FK2 antibody, in 5% BSA in PBS-Tween20, and incubated with primary antibodies overnight at 4°C, followed by horseradish peroxidase-conjugated anti-mouse (Cell Signaling) or anti-rabbit (Cell Signaling) secondary antibodies for 60 min at RT. Blots were developed with Super Signal West Pico ECL (Thermo Fisher Scientific). Blots were imaged using Amersham Imager 600 chemiluminescence imager (GE Healthcare Life Sciences). Primary antibodies used for Western blotting and their dilutions are listed in the Reagents and Resources table. Densitometric evaluations of protein expression were performed using the ImageJ64 image analysis software (NIH, Bethesda, MD), as reported ^21,54^.

### MYC- and GFP-immunoprecipitation

Immunoprecipitation of MYC-tagged proteins was done as described^21,3^. Cells were collected and suspended ice-cold IP buffer (20 mM Tris-HCl, pH 7.5, 150 mM NaCl, 1 mM EDTA, 0.5% NP-40, 5 mM *N*-ethylmaleimide, and protease inhibitors), followed by incubation on rotator for 1 hr at 4°C. Samples were spin down at 20,000 x g, at 4°C for 45 min, and supernatants containing solubilized material were transferred to new tubes. Proteins were immunoprecipitated with anti-MYC mAb-conjugated agarose beads (Clontech Takara), washed 4 times in IP buffer, and eluted using 2.5% acetic acid. For GFP-immunoprecipitation, samples prepared as above were immunoprecipitated using agarose beads-conjugated anti-GFP nanobodies (GFP-TRAP; Protein Technology). To achieve removal of co-immunoprecipitating proteins, samples were washed 3 x 20 min at RT with IP buffers supplemented with 1%SDS, and then 2 x 20 min with regular IP buffer. Samples were eluted by incubation with 2 x SDS PAGE sample buffer for 10 min at 95°C.

### Immunofluorescence

Immunofluorescence was performed as previously described ^3,21,38^. Briefly, cells grown in 2- or 4-well chamber slides (model 1 German borosilicate; Lab-Tek; VWR) were fixed with freshly prepared 4% formaldehyde in PBS solution (using 16% Methanol-free Formaldehyde; Thermo Fisher Scientific) for 20 min at RT, then permeabilized with permeabilization buffer (PB; 0.15% Triton X-100 in PBS) for 20 min at RT. After blocking with blocking buffer (BB; PB supplemented with 7.5% BSA) for 45 min, samples were incubated with primary antibodies suspended in BB for 90 min at RT, followed by 3 washes with BB and incubation with secondary antibodies diluted in BB for 60 min at RT. Primary antibodies and their dilutions are listed in the Reagents and Resources table. Secondary antibodies were highly cross-absorbed goat anti-mouse Alexa Fluor-488 and goat anti-rabbit Alexa Fluor-546 (1:1000; both from Thermo Fisher Scientific). In triple labelling experiments, cells were immunostained, as above, followed by 3 washes with BB and incubation with anti-Tom20 antibodies conjugated with Alexa 647 fluorophore (Santa Cruz Biotechnology Inc.; 1:100) for 90 min at RT. After 3 washes with PBS, cells were subjected to Airyscan imaging. Immunolabeled cells on chamber slides were stored in PBS at 4°C and imaged within 10 days after processing.

### Image acquisition, analysis and processing

Images were acquired with a Zeiss LSM 880 microscope (Zeiss MicroImaging) equipped with an Airyscan superresolution imaging module, using 63/1.40 Plan-Apochromat Oil DIC M27 objective lens (Zeiss MicroImaging), as described ^3, 21,54^. The 488-nm Argon laser line, 561-nm DPSS 561 laser, and 633-nm HeNe 633 laser were used to detect Alexa-488, Alexa-546, and Alexa-647, respectively. The *z*-stacks covering the entire depth of cells with intervals of 0.18 μm were acquired. Image processing and analyses were done using ZEN software (version 3.9, Zeiss MicroImaging) and ImageJ (NIH, Bethesda, MD). Prior to analyses, all images were processed using Joint Deconvolution, resulting in generation of 90-nm lateral resolution images, as described^3^. For peroxisome number and size, single cells in maximum-intensity projection images were outlined, cropped, and converted into binary images using ImageJ, and analyzed using the “particle analysis” module of the software^3^. The data were tabularized and transferred to Microsoft Excel software (Microsoft) for further analyses. Image cropping and global brightness and contrast adjustments were performed for presentation using Adobe Photoshop CS6 software (Adobe Systems).

### Mass spectrometry-based proteomics

Cell lysis and protein digestion were carried out as previously described ^55–57^. Briefly, samples were lysed in a buffer composed of 5% sodium dodecyl sulfate (SDS) and 50 mM triethylammonium bicarbonate (TEAB; 1 M, pH 8.0). Protein samples were sonicated using a probe for 30 s. Protein concentrations were quantified using a BCA assay (Cat#). Lysates were subsequently digested using S-Trap micro columns (ProtiFi, NY) according to the manufacturer’s instructions. Peptides were eluted, dried, and reconstituted in 0.1% formic acid. Peptide concentrations were determined using a BCA protein assay kit (Thermo Fisher Scientific, 23275). All tryptic peptides were separated on a nanoACQUITY UPLC analytical column (BEH130 C18, 1.7 µm, 75 µm × 200 mm; Waters Corporation, Milford, MA) using a 120-min linear gradient of 3–50% acetonitrile with 0.1% formic acid, operated on a nanoACQUITY UPLC system (Waters Corporation). The eluate was directly analyzed on an Orbiztrap Fusion Lumos Tribrid mass spectrometer (Thermo Scientific, San Jose, CA) operating in data-independent acquisition (DIA) mode. DIA settings were as follows: MS1 scan range, 400–850 m/z; MS2 scan range, 145–1450 m/z; isolation window, 3.0 m/z; HCD collision energy, 28%; Orbitrap resolution, 6000; and maximum injection time, 54 ms.

DIA raw data were processed using DIA-NN software (version 2.2.0) ^58^. Searches were performed against the *Homo sapiens* UniProtKB/Swiss-Prot database (downloaded March 2025; 20,404 entries) using the library-free workflow. The following options were enabled: *FASTA digest for library-free search/library generation* and *Deep learning spectra, RTs and IMs prediction*. Search parameters included a maximum of one missed cleavage, up to three variable modifications, and peptide length range of 6–50 amino acids. Carbamidomethylation (Cys) was set as a fixed modification, while N-terminal methionine excision was included as variable modification. The precursor charge range was set to 2–6, precursor m/z range to 300–1000, and fragment ion m/z range to 145–1450. Precursor and fragment ion mass tolerances were set to 8 ppm and 20 ppm, respectively. False discovery rates (FDRs) at the protein and peptide levels were controlled at 1%. Match-between-runs was enabled. The resulting pg.matrix file containing protein abundance values was analyzed using Perseus (version 1.6.14.0). To ensure robust statistical analysis, proteins with missing values across any biological replicates were excluded. Quantitative protein intensities were log_₂_-transformed and normalized by median centering. Differentially expressed proteins were determined using two-tailed Student’s *t*-test, with adjusted *p* < 0.05 considered statistically significant. Ingenuity Pathway Analysis (IPA) (Qiagen, Germantown, MD) was used to identify affected pathways and biological processes.

### Key resources table

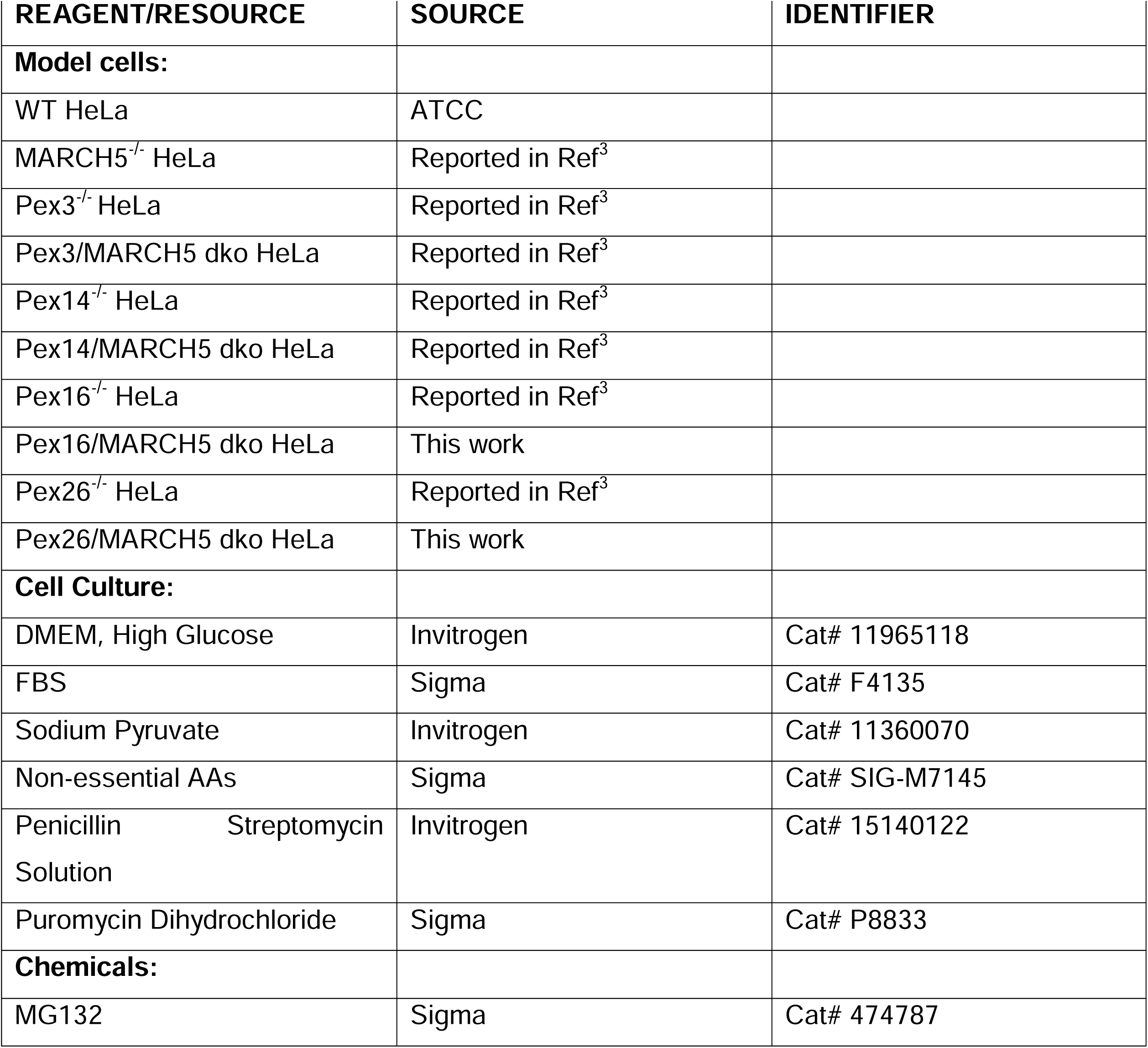
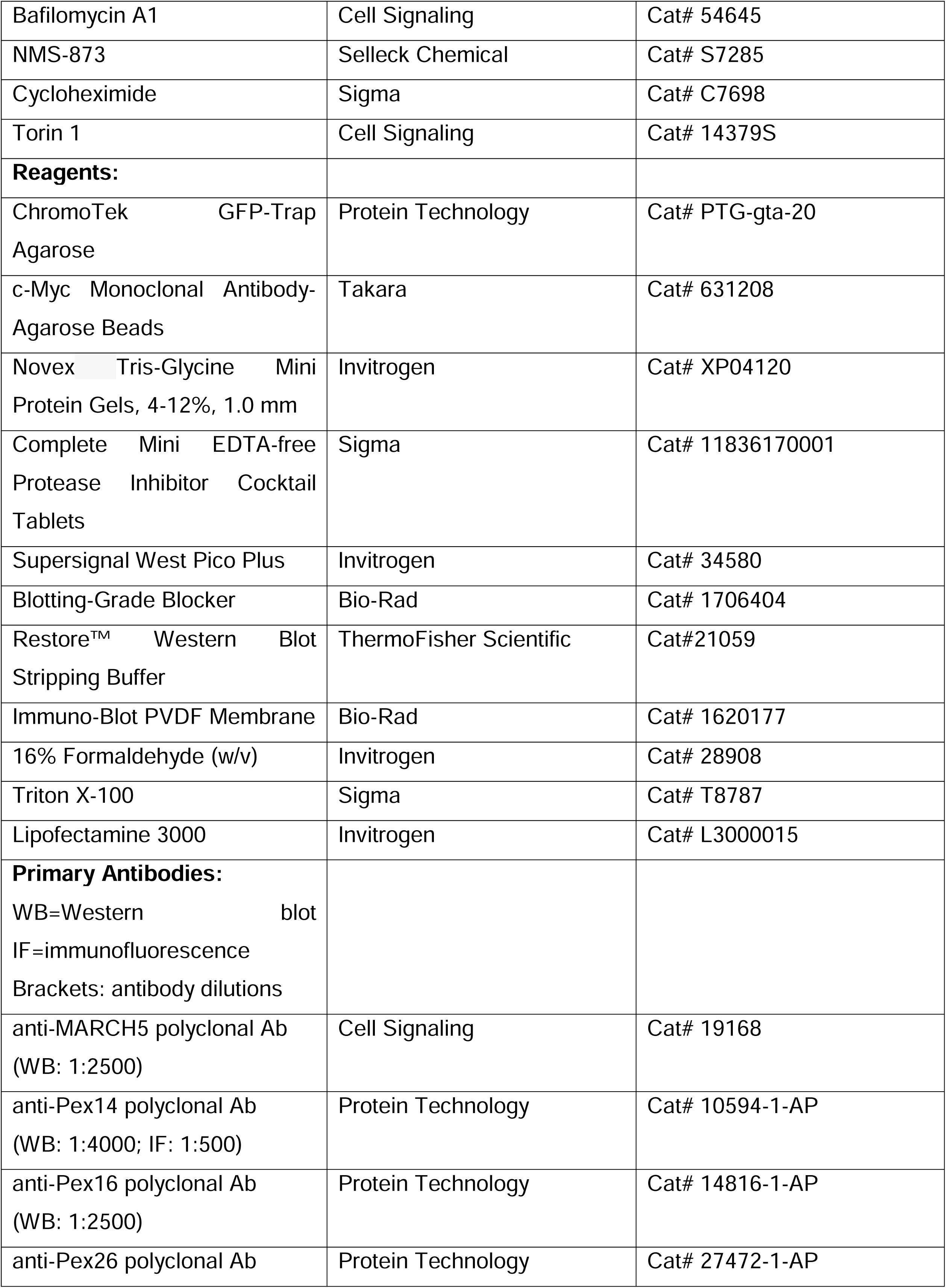
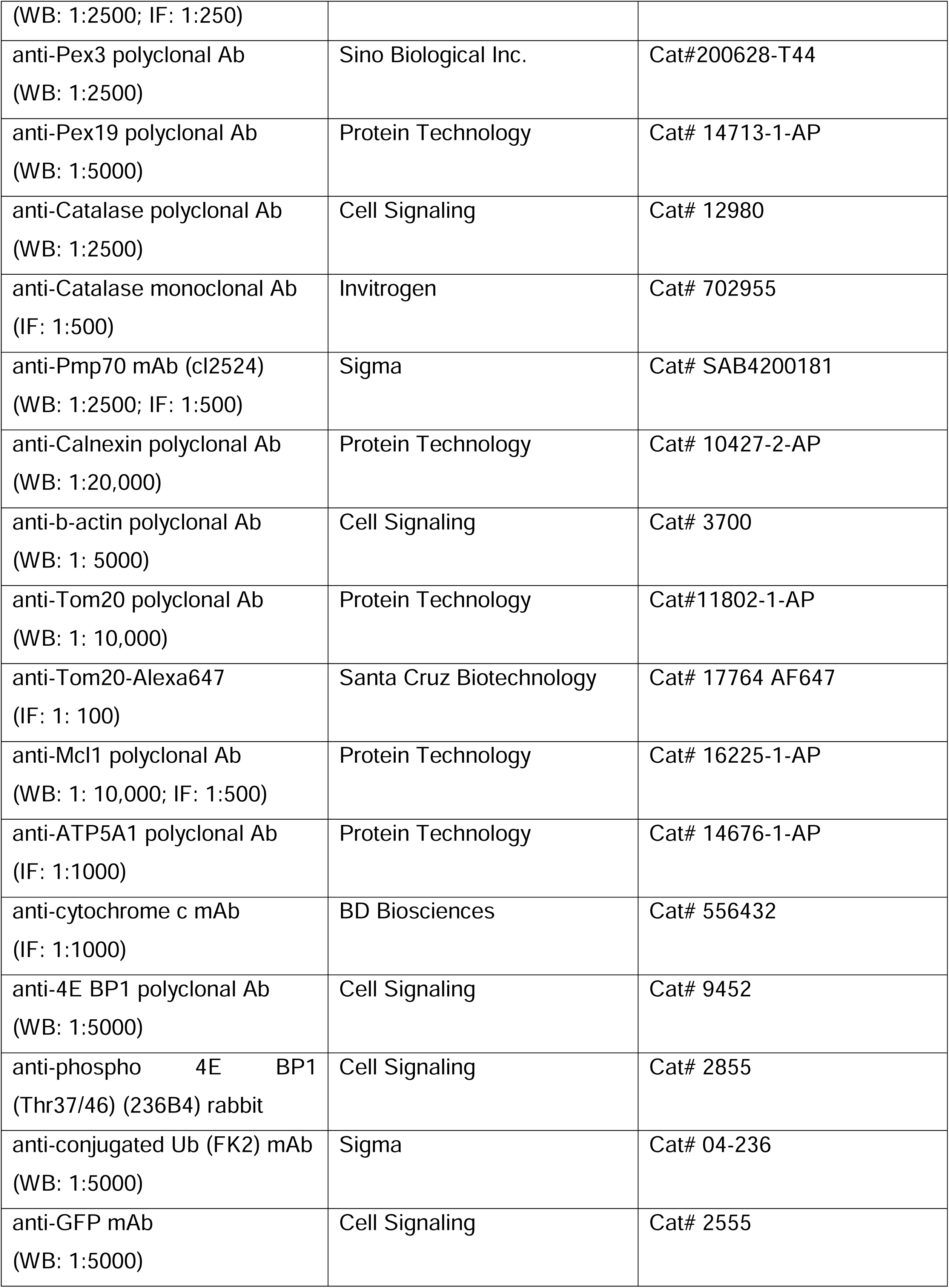
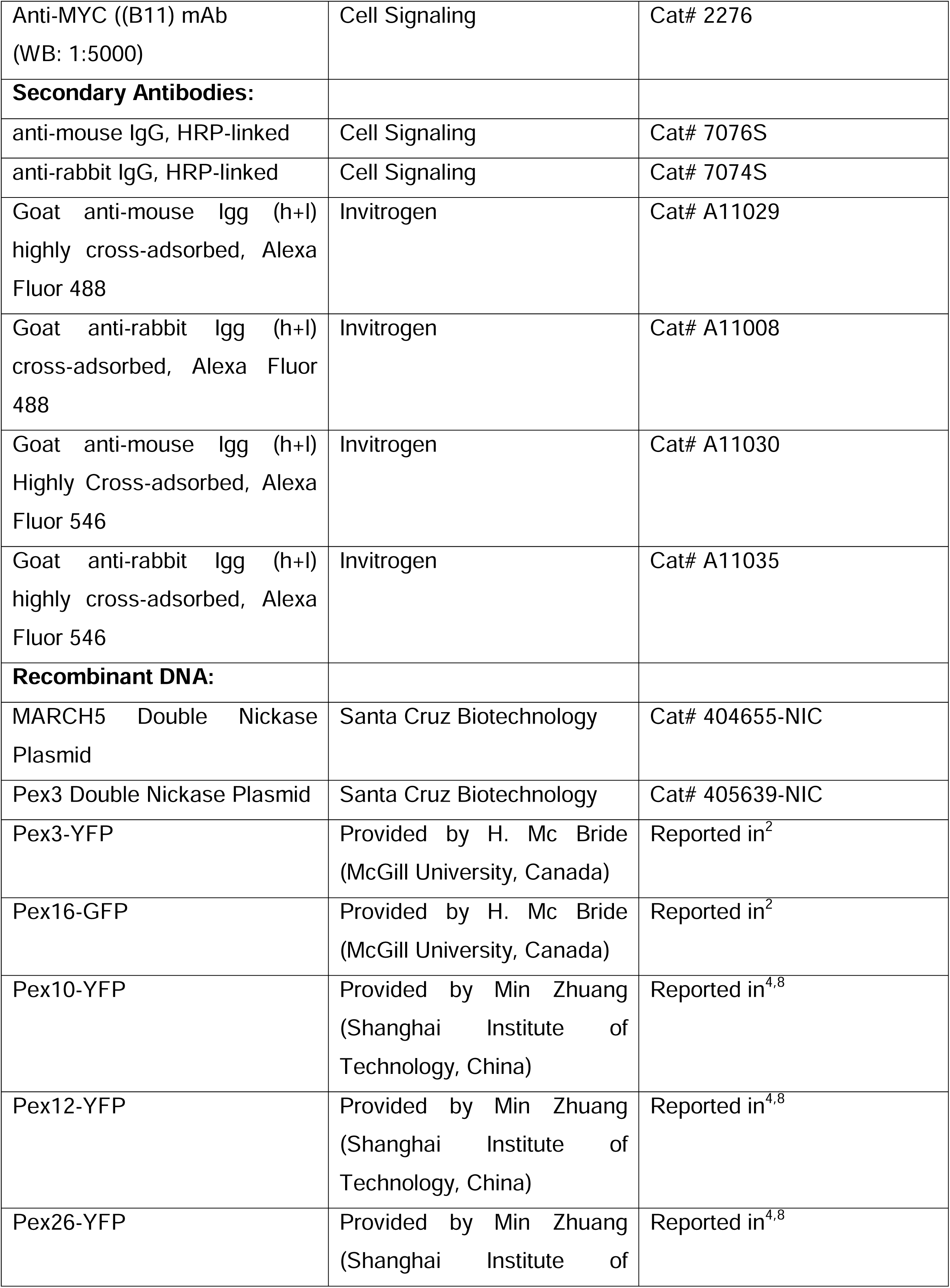
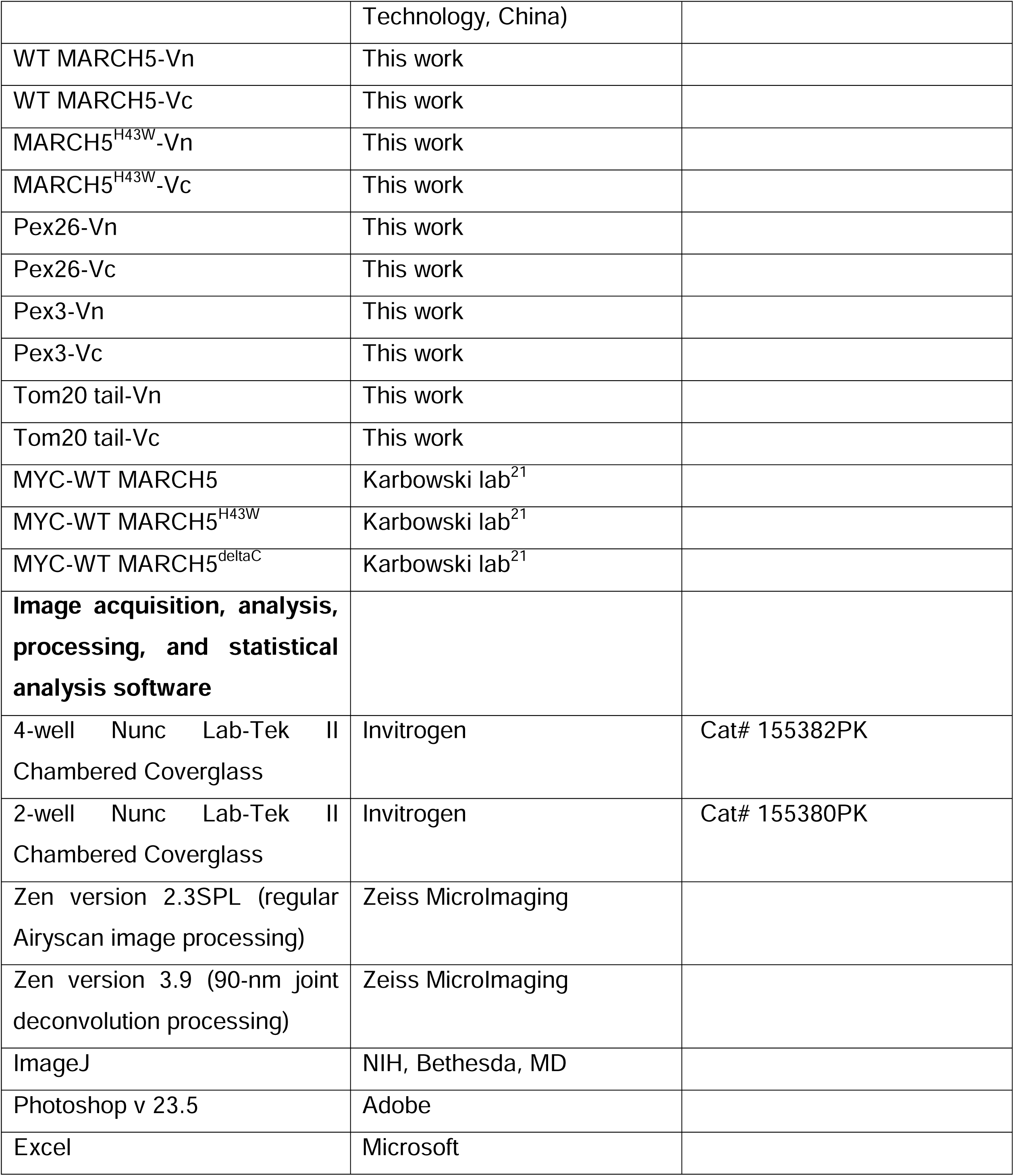

## Acknowledgements

We thank Nicolas Verhoevem for help in earlier stages of this project, Wieslawa Jarmuszkiewicz, Wojciech Pijanowski, Krzysztof Wojcicki, and Michael Wagner for critical reading of the first draft of the manuscript. Research reported in this publication was supported by the National Institutes of Health under Award Number 1R01GM153862-01A1 (MK) and by the Center for Biomedical Engineering and Technology (BioMET), University of Maryland, Baltimore.

## Author contribution

DB and MK conducted the experiments and analyzed data, CCB and AN generated critical research tools, GZ provided access to imaging resources, LB performed all statistical analysis, and MMW and MAK designed, performed, and analyzed proteomic studies. MK designed the study, supervised the project, and wrote the manuscript. All authors read and influenced the current version of the manuscript.

## Supplemental Figure Legend

**Figure S1 (related to Figure 1A):**
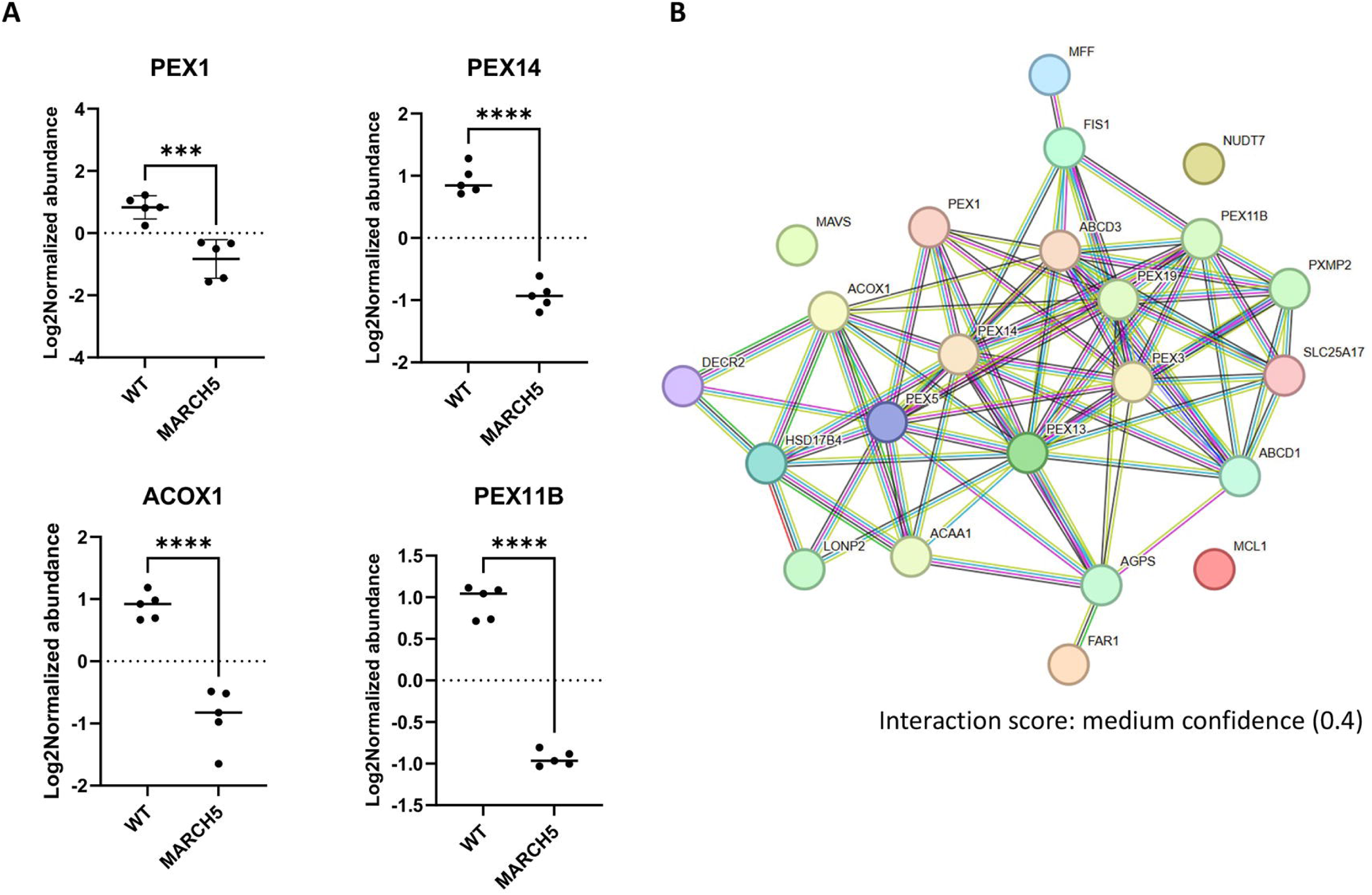
(**A**) Representative changes in protein expression within the peroxisome import pathway. (**B**) Protein–protein interaction network analysis of peroxisome-associated proteins in MARCH5^−/−^ compared to WT.

**Figure S2 (related to Figures 2 and 3):**
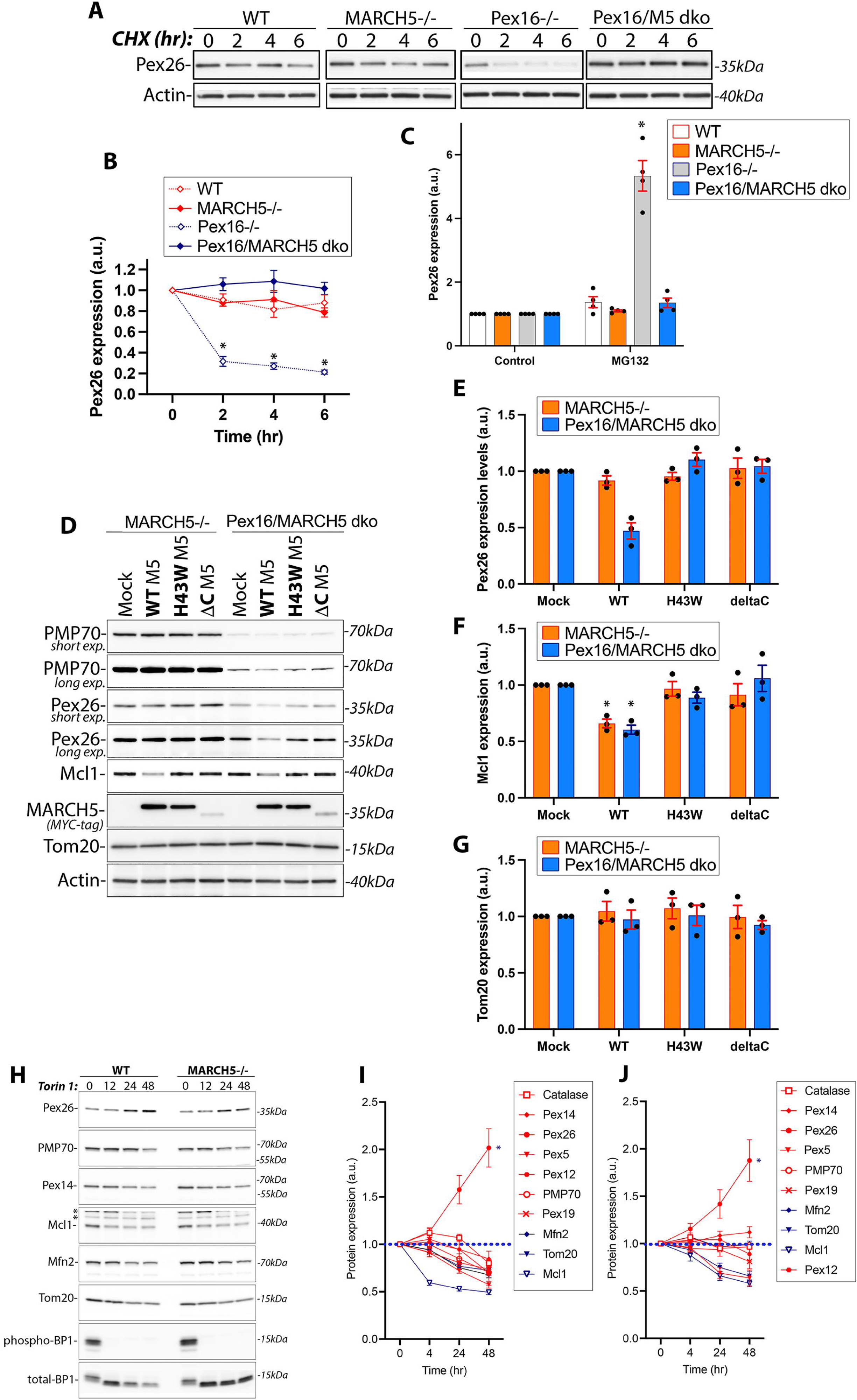
Effect of MARCH5 deficiency on Pex26 turnover in Pex16^−/−^ cells. **(A)** Cells indicated in the figure were treated with CHX for 0-6 hr, followed by Western blot to detect Pex26. Actin is a loading control. **(B)** Changes in Pex26 expression levels in cells treated as in **A** were quantified. Value at 0 hr was taken as 1. Mean ± SEM. N=3. Unpaired t-tests and Wilcoxon rank-sum tests. Multiple comparisons were corrected using the Benjamini-Hochberg FDR method. (* p<0.05) **(C)** Cells indicated in the figure were treated with MG132 for 6 hr, followed by densitometry and quantification of levels of Pex26. Values in untreated cells were taken as 1. Mean ± SEM. N=4. **(D)** Western blot analyses of total cell lysates from MARCH5^−/−^ and Pex16/MARCH5 dko cells, mock transfected, or transfected with MYC-WT MARCH5 (WT M5), MYC-MARCH5^H43W^ (H43WM5) and MARCH5^deltaC^ (ΔCM5). Tom20 and Actin are loading controls. (**E**-**G**) Quantification of expression levels of Pex26 (**E**), Mcl1 (**F**), and Tom20 (**G**) in cells treated and analyzed as in **D**. Mean ± SEM. N=3. Levels of respective proteins in ‘mock” transfected cells were taken as 1. Each phenotype was compared to “mock” using unpaired t-tests and Wilcoxon rank-sum tests. Multiple comparisons were corrected using the Benjamini-Hochberg FDR method. (* p<0.05) **(I)** Western blot analyses of total cell lysates from WT (left panels) and MARCH5^−/−^ (right panels) HeLa cells, treated with Torin1 for 0, 4, 24 and 48 hr. Peroxisomal and mitochondrial proteins were analyzed as indicated in the figure. Reduction of phosphorylated 4E-BP1 (eukaryotic translation initiation factor 4E-binding protein 1) was used as a control for Torin1 efficacy in both cell types. (**I**, **J**) Quantifications of the expression levels of the indicated proteins in WT (**I**) and MARCH5^−/−^ (**J**) Hela cells treated as in **H**. N=3-6 independent experiments. Peroxisomal proteins are in red, mitochondrial in blue. Mean ± SEM. N=4-7. Statistics (**I**-**J**): paired t-tests comparing each later timepoint to 0 h, with Benjamini-Hochberg FDR correction across proteins. Stars denote FDR-corrected significance (* p<0.05, ** p<0.01, *** p<0.001). (**I**) *-catalase, Pex26; **-Pex19, PMP70, Mcl1, Tom20: ***-Mfn2, Pex5, (**J**) *-Pex26, Mcl1, Pex5, Tom20.

**Figure S3 (related to Figures 3 and 4):**
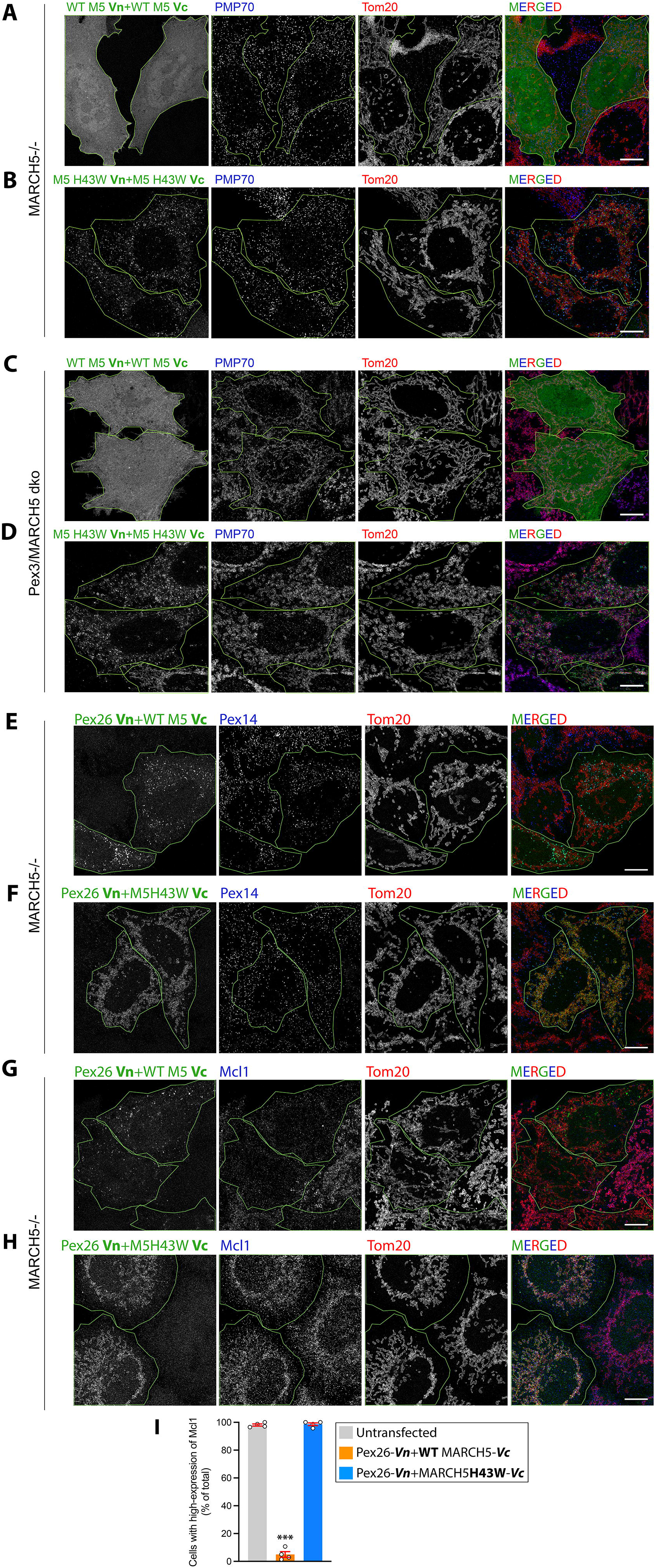
BiFC control experiments. (**A**-**H**) MARCH5^−/−^ (**A**,**B**,**E**-**H**) and Pex3/MARCH5 dko (**C**,**D**) transfected with BiFC constructs indicated in the figure (green on merged images) were immunostained to detect peroxisomal proteins PMP70 (**A**-**D**; blue), and Pex14 (**E**,**F**; blue), or MARCH5 client protein Mcl1 (**G**, **H**; blue) together with the OMM marker (Tom20, **A**-**H**; red). Bars:10μm. **(I)** Quantification of Mcl1 expression levels in MARCH5^−/−^ cells transfected with Pex26-Vn+WT MARCH5-Vc (as in **G**), or Pex26-Vn+MARCH5^H43W^-Vc (as in **H**). Mean ± SEM. N=4 (>50 cells counted each time). Cells showing significantly reduced Mcl1 signal (as in **G**) were considered to have low Mcl1 expression levels. Statistics (**I**): Each phenotype was compared to WT using unpaired t-tests and Wilcoxon rank-sum tests. Multiple comparisons were corrected using the Benjamini-Hochberg FDR method.

